# Computational Insights into Colonic Motility: Mechanical Role of Mucus in Homeostasis and Inflammation

**DOI:** 10.1101/2023.08.21.554097

**Authors:** I.H. Erbay, A. Alexiadis, Y. Rochev

## Abstract

Colonic motility plays a vital role in maintaining proper digestive function. The rhythmic contractions and relaxations facilitate various types of motor functions that generate both propulsive and non-propulsive motility modes which in turn generate shear stresses on the epithelial surface. However, the interplay between colonic mucus, shear stress, and epithelium remains poorly characterized. Here, we present a colonic computational model that describes the potential roles of mucus and shear stress in both homeostasis and ulcerative colitis (UC). Our model integrates several key features, including the properties of the mucus bilayer and lumen contents, colonic pressure, and crypt characteristics to predict the time-space mosaic of shear stress. We show that the mucus thickness which could vary based on the severity of UC, may significantly reduce the amount of shear stress applied to the colonic crypts and effect colonic content velocity. Our model also reveals an important spatial shear stress variance in homeostatic colonic crypts that suggests shear stress may have a modulatory role in epithelial cell migration, differentiation, apoptosis, and immune surveillance. Together, our study uncovers the rather neglected roles of mucus and shear stress in intestinal cellular processes during homeostasis and inflammation.

## 1. Introduction

Intestinal mucus covers the surface of the intestinal epithelium and acts as a biochemical defence barrier, lubricant, and selective transport system (1, 2). The properties and organisation of human intestinal mucus are site-dependent to address specific needs (3). In the colon, mucus consists of a stratified inner layer (SL) and a loose outer layer (LL) which together forms a bilayer (4). The SL thought to protect the underlying epithelium from mechanical forces and maintains sterility, while the LL is used as a food source for the microbiome and as a lubricant for faeces (5, 6). The current *in vitro* and computational studies of intestinal mucus mainly focus on adhesion/penetration characteristics for various nanoparticle formulations, glycosylation profiles in homeostasis and disease, and secretion mechanisms (7-9). Further, a large portion of mechanistic studies for the epithelial barrier is studied either under static conditions or without the mucus layer. In advanced cell models such as organoids and induced pluripotent stem cells (iPSCs) and their combinations with organ-on-a-chip (OoC), the mucus layer is either missing as a layer and shown as a protein expression in the models or fail to form a distinct mucosal layer. Consequently, although the importance of shear stress has been repeatedly shown in OoC models, no general framework exists to account for the interplay of shear stress, epithelial barrier, and intestinal mucus (10-18).

To sustain homeostasis and withstand repetitive forces, including shear stress caused by luminal flow, stretching, and compression of the tissue wall during transport, mixing, and absorption of lumen contents, the epithelial barrier is continuously regenerated (19, 20). One of the critical roles of colonic mucus is to protect the epithelium from mechanical forces during faecal propulsion and intercellular forces caused by the extrusion of adjacent cells during cell migration from the crypt base towards the lumen (21). Neural peristalsis is the main propulsive event that can affect approximately 30 cm of the colon at a time and is characterized by various rhythmic waves (22). During peristalsis, intraluminal pressure increases significantly and induces segmental motility while generating shear stress on the epithelial barrier surface (23-26). Despite studies on the regulation and functional pathways of stem cells and mechanical analysis of intercellular contact providing insights into complex cellular interactions, the variance in mechanisms during inflammation is not well understood (27, 28). Moreover, the role of spatial shear stress on intestinal crypts and mucus, which may be significant in homeostatic processes, including cell migration, cell shedding, immune cell recruitment, and pathogenic pathways, is often disregarded. For instance, imbalance in the mucus barrier reportedly plays an important role in the progression of inflammatory bowel disease (IBD) and colorectal cancer (29-31). Furthermore, the thickness and density of the mucus layer are decreased in active ulcerative colitis (UC), potentially leading to increased bacterial translocation and immune activation (32, 33).

To date, computational models of intestinal processes have been used to characterize and model the fluid mechanics that drive digestion and nutrient absorption. Among various numerical techniques, mesh-based and particle-based methods such as discrete multiphysics (34) are frequently used to model intestinal flow (35, 36), mixing (37), and absorption (38) in humans. However, the mucus layer is often overlooked, and its contribution to these processes is crucial, as dysregulation often leads to serious conditions. Developing a better understanding of intestinal processes and building close-mimicry systems requires a deeper insight into the role of mucus during intestinal motility.

Here, we present a computational model that describes the time-space interplay between the mucus layer and shear stress in colonic crypts during a peristalsis-like motion. Our approach is based on a three-phase field and fluid-solid interaction method to represent the basic biomechanical microenvironment. We used experimental data on colonic pressure, mucus rheology, and crypt characteristics to predict shear distribution in homeostasis and UC. Our models include a variation in mucus thickness from 0 to 800 µm to account for severity of UC, during which mucus thickness decreases. By incorporating key components, such as the mucus layer, colonic contents, and epithelium, our model provides important insights into the time-space mosaic of mechanical forces, as well as how mucus thickness affects the spatial shear stress distribution. Our work opens up exciting prospects for studying epithelial barrier processes and provides a framework for mechanistic organ-on-a-chip models to develop novel pharmaceutical targets.

## 2. Methods

### 2.1 Geometry construction

The human colon consists of small pouches collectively called haustra, although the size of the haustrum varies throughout the colon, each has approximately a length of 30 mm and a radius of 50 mm. In contrast to the small intestine, villus structures are missing in colonic crypts, which form the intestinal epithelium. The geometrical parameters of the human colon, including height, radius, and distance between two crypts, were adapted from previously published studies, and an average value was calculated for each feature. The geometric parameters are listed in Table 1. A 2D rectangular representation was then constructed to reduce the computational time. The crypt features were assumed to be uniform throughout the haustrum. Colonic mucus was constructed as a bilayer, low-viscosity loose layer (LL), and higher-viscosity stratified layer (SL) that separates the crypts from the lumen. The thickness of the colonic mucus varies depending on the site; moreover, the reported thicknesses contradict each other varying between 100-800 microns. In this study, we investigated various thicknesses of SL mucus, including 100, 200, 400, 600, and 800 µm. Furthermore, the lumen was assumed to be filled with faeces. The physical parameters of the mucus and colonic contents are listed in Table 2.

**Table 1.** Geometrical parameters of human transverse colon.

| Parameter | Value | Reference |
| --- | --- | --- |
| Length of the haustrum | 30 mm | (26) |
| Height of the haustrum | 50 mm | (39) |
| Height of a crypt | 400 $\mu\text{m}$ | (40) |
| Radius of a crypt | 50 $\mu\text{m}$ | |
| Distance between two adjacent crypts | 200 $\mu\text{m}$ | |
| Length of a crypt | 30 mm |  |
| Thickness of the LL | 100 $\mu\text{m}$ | (41, 42) |
| Thickness of the SL | 100 – 800 $\mu\text{m}$ | |
| Length of mucus layers | 30 mm |  |
| Crypt density per $\text{mm}^2$ | 14 $\text{mm}^{-2}$ | (40) |

**Table 2.** Properties of faeces and mucus bilayer.

| Carreau fluid parameters |  |  |  |  | Power law fluid parameters |  |  |
| --- | --- | --- | --- | --- | --- | --- | --- |
| Component | Zero shear rate viscosity [Pa s] | Infinite shear rate viscosity [Pa s] | Power index (n) | Relaxation time ($\lambda$) [s] | Fluid consistency coefficient [Pa s] | Flow behaviour index (n) | Density [kg m<sup>-3</sup>] |
| LL | 0.1 | 0 | 0.375 | 0.001 | - | - | 1000 |
| SL | 21 | 0.1 | 0.375 | 15 | - | - | 1400 |
| Faeces | - | - | - | - | 682 | 0.31 | 1060 |

### 2.2 Governing equations

A finite element based commercial computational software, COMSOL Multiphysics® (Version 6.1, COMSOL Inc., Burlington, MA), was used to develop the numerical model. Two groups of simulations were carried out, first a model that consisted of only lumen and the crypt structure, second a model with lumen, mucus bilayer with various SL thicknesses, and crypt structures. For the first group, a Fluid-Structure Interaction (FSI) model with single-phase laminar flow was employed. For the second group, simulations were based on a ternary-phase field model for colonic contents, and mucus layers to study the evolution of three fluid phases denoted by fluid A, fluid B and fluid C. The fluid phases were faeces, LL, and SL respectively. We considered both mucus layers to be in reasonable agreement with Carreau fluid (43) and faeces as a power law fluid (44).

#### 2.2.1 Fluid domain

The fluid domain consisted of laminar flow and ternary phase-field models. The continuity and momentum equations for laminar flow are given, respectively, by.

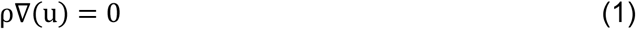

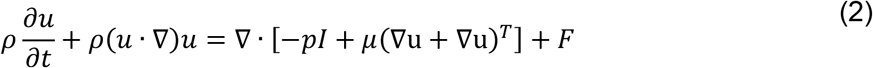

where ρ is the density of the fluid [kg m^-3^]; *u* is the fluid velocity [m s^-1^]; *p* is pressure [Pa]; **I** is the identity matrix; *μ* is the dynamic viscosity of the fluid [Pa s]; and **F** is the volume force vector describing a distributed force field [N m^-3^].

The ternary model consists of LL, SL, and faeces. A ternary field model was used to study the evolution of the phases (45). The model solves the following Cahn-Hillard equations assuming the phases are immiscible:

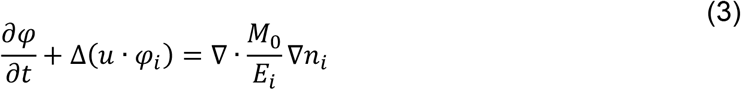

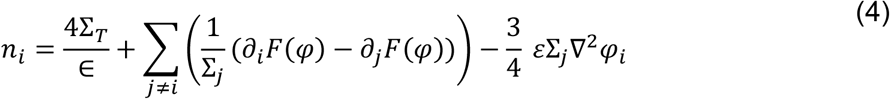

governing the phase field variable, *i*, and a chemical potential, *n*_*i*,_ for each phase *i*=A, B, C.

where *ε* is the parameter determining the thickness of the interface; *M*_*0*_ is a molecular mobility parameter and *Σ*_*T*_ is defined as:

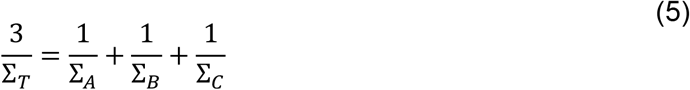

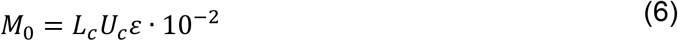

Where *L*_*c*_is the characteristic length and *U*_*c*_is the velocity scale of the system.

The phase field variables vary between 0 and 1. At each point the phase field variables satisfied the following condition:

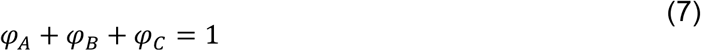

The density of each phase is assumed to be constant, that is the phase field variable is equivalent to the volume fraction of the phase being considered. The density and viscosity of the fluid mixture applied in (1) and (2) are defined as:

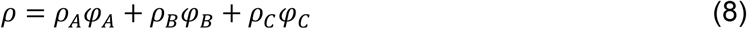

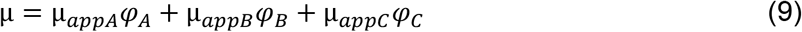

Both SL and LL were considered to be incompressible shear-thinning non-Newtonian fluids. The Carreau model was used to describe the dynamic viscosity *μ* as a function of the strain rate γ^·^ as follows:

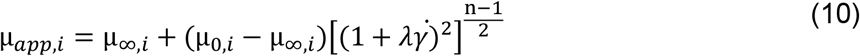

Where *µ*_*∞i*_ is the viscosity of phase *i* at infinite shear, *µ*_*0,i*_ is the viscosity of phase i at zero shear, *λ* is characteristic time constant, γ^·^ is shear rate, and n is power law index.

Power law

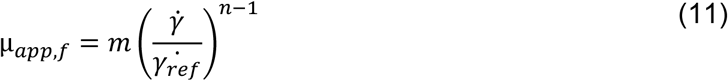

The free energy of the 3-phase system is defined as a function of the phase field variables as:

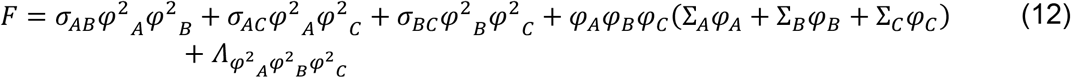

Where *σ*_*AB*_ denotes the surface tension coefficient of the interface, separating phase A and B; is the parameter specifying the additional free bulk energy. The capillary parameters *Σ*_i_ are defined for each phase as follows:

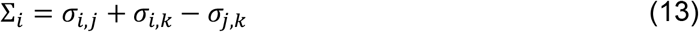

The surface tension force applied as a body force, was calculated from the chemical potentials as follows:

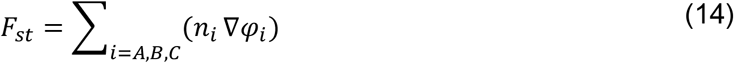

Shear stress was evaluated by multiplying shear rate and dynamic viscosity as follows:

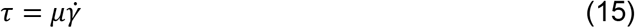

#### 2.2.2. Solid Domain

FSI modelling is used to study the interaction between fluid and solid structures, which occurs at their boundaries and interfaces. The fluid forces acting on the solid walls in a flowing stream create stress and deformation in the solid body. In the FSI, the problem is solved for stresses and deformations in the deformed solid body, as well as for the velocities and pressures in the fluid flow. The deformation of the structure influences the flow variables in the fluid, and it can be small or large, depending on the material properties, pressure, and velocity. As the stiffness of the tissue significantly increased under UC conditions, we reflected this in the material properties, which are given in Table 3. To handle the FSI, a non-slip boundary was set on the top layer of the crypt structure, embedded in the SL, using the arbitrary Lagrangian-Eulerian (ALE) approach. A time-dependent, fully coupled approach was used to define coupling along the FSI boundary under no-slip boundary conditions.

**Table 3.**
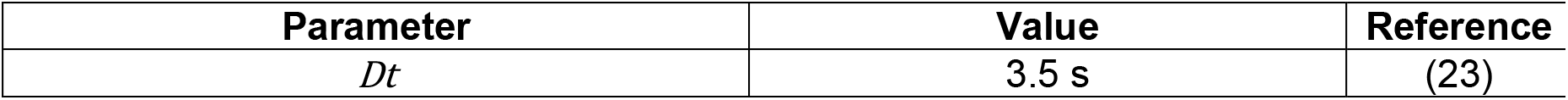
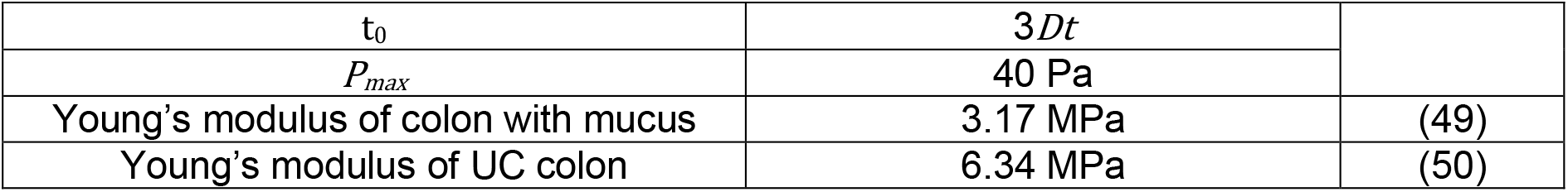
Motility and mechanical parameters.

A moving mesh model was used to study time-dependent deformations and geometrical changes due to the motion of boundaries in FSI (46). These conditions can be expressed as:

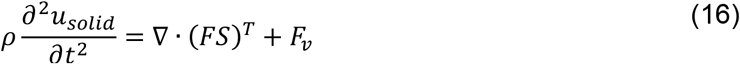

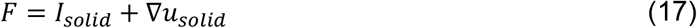

where *u*_*solid*_ is the displacement in the solid body, *ρ* is the body’s structural density, **S** is the 2nd Piola-Kirchhoff stress tensor, **F** is the deformation gradient, and **F**_**v**_ is the body force, **I**_***solid***_ is the identity matrix.

A fully coupled solver used with time stepping determined automatically using a backward differentiation formula. Finally, hydrodynamic shear stress was calculated as the product of shear rate and fluid viscosity.

### 2.3 Boundary conditions

The crypt structure was fixed at the base and top points while being impermeable to fluid flow. No-slip boundary conditions were defined between the mucus layers and the crypt domain. The inlet (*x*=0 mm) and outlet (*x*=30 mm) boundary conditions were driven by the applied pressure, and the inlet flow was calculated using Eq. 18 The outlet pressure was set as 0 [Pa]. A physics-controlled adaptive mesh strategy was employed to ensure the sufficient fineness of the computational meshes used in the simulations. All simulations were completed with a time-dependent study, in which the duration ranged from 0 to 20 [s].

### 2.4 Intestinal motility specification

Intestinal motility includes complex patterns of either propulsive or non-propulsive motor activities that may occur in both the oral and aboral directions. The primary propulsive activity is peristalsis, which results from the pressure waves generated by the muscles. To generate a peristalsis-like propulsive wave in our model, we used pan-colonic manometry data from healthy subjects (23-25). Simultaneous pressure waves (SPWs) describe superimposed pressure peaks on the downward slope. We simplified the SPWs to fit a Gaussian curve to expand the range of pressures in the system using the following equation:

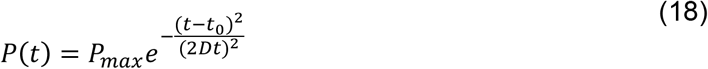

where *P*(*t*) is the pressure, *P*_*max*_ is the peak value of the pressure, *Dt* is the standard deviation. Pressure is applied to the inlet to generate a hypothetical peristalsis-like push in the system. Because the geometry can deform, this approximates the physiological conditions. However, as the pressure values in these studies reflect the total pressure increase rather than the active pressure that causes motility, we adjusted the inlet pressure based on MRI-based velocity measurements and the total mass accumulation at the outlet (47, 48).

### 2.5 Statistics

Statistical analysis was completed by selecting the same boundaries on crypts at the top and bottom of the lumen in every simulation (n=6) consisting of UC model without mucus and models with 100, 200, 400, 600, and 800 µm of SL thicknesses. The selection was carried out every 3 mm from 7 mm to 27 mm. The shear stresses on these points were then taken as averages for all time points. All results are presented as mean ±standard deviation (SD). For the direct comparison of six groups consisting of models, two-way analysis of variance (ANOVA) with Tukey’s test was used to compare the means of all groups. *P<0.05; **P<0.01; ***P<0.001; ****P<0.0001; n.s., not significant (P>0.05).

## 3. Results and Discussion

### 3.1 Haustral geometry construction and pressure wave guided flow condition

The colon consists of series of haustrum, each approximately 5 cm in length and 2.5 cm in diameter. Using histological sections from the literature (43, 44, 51), we identified the height, length, radius, and distance between two adjacent crypts. Furthermore, a mucus bilayer was constructed on top of the crypts as a barrier to the lumen. SL was built on top of the crypts, and LL was built on top of SL. The thickness of the human colonic mucus is a point of debate and reported values span from 100 to 800 µm, which can be traced back to the challenges in measuring it under physiological conditions (3, 29, 41). Furthermore, a decrease in mucus thickness is a typical pathology of UC and depends on the severity of the condition (32). Consequently, we varied the thickness of the SL to 100, 200, 400, 600, and 800 µm while maintaining the LL at 100 µm. The height of the lumen naturally decreases as the thickness of the mucus layer increases. This allowed us to simulate not only severe UC conditions, where there is no mucus but also milder pathologies, where the mucus thickness is reduced. We simplified the haustrum geometry as a rectangle containing colonic crypt structures on both sides of the surface. Colonic crypt length, width, and distance between two crypts can vary depending on the colonic site and age of the crypt. We assumed that crypt geometry was uniform throughout the model (Fig. 1a). In order to understand how flow affects the crypt structures, we identified three biologically relevant sites in our models. Namely, crypt base where the stem cells reside, crypt migration route which is the length along the crypt from base to top where cells start to differentiate into either secretory or absorptive cell types, and finally crypt top where cells are matured and undergo apoptosis after their life-cycle.

**Figure 1.**
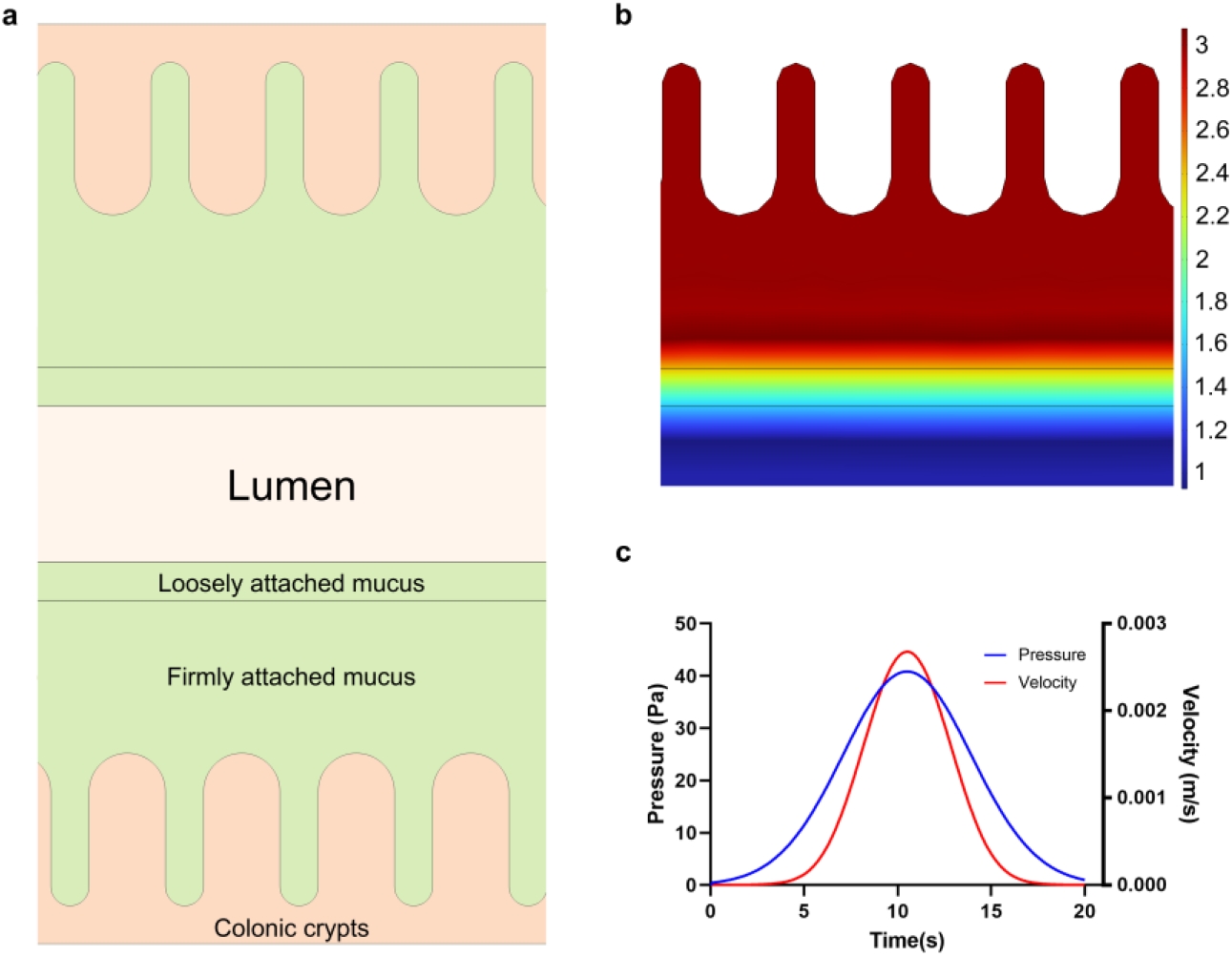
Schematic diagram of the simplified human haustrum geometry, pressure wave, and interfaces between phases: (a) 2D representation of the lumen with colonic crypt structures distributed on the inner surface, protected by the mucus bi-layer (b) Interface of faeces, loosely attached mucus (LL), and firmly attached mucus (SL) at the start of the simulation (t=0). The colour bar indicates the purity of each phase (3: Firmly attached mucus, 2: Loosely attached mucus, and 1: Faeces (c) Line graph of the pressure wave (Pa) and velocity (m/s) over time at the inlet to induce a peristalsis-like motion in the system.

To model the mucus layers, we employed a ternary-phase field method and combined it with FSI to model the mucus bilayer, faeces, and subsequent interactions between the flow and crypt structures. Ideally, the motility of the mucus layers will be minimal, while the pressure wave transports the faecal phase in the system (Fig. 1b). The pressure in the system was introduced as a function of maximum pressure (40 Pa) and time, and the velocity in the system follows a similar distribution, reaching approximately the maximum values at approximately t=10s in the total simulation of 20s (Fig. 1c). The movement of colonic contents is associated with measurable pressure events that present regional variance in quantity and characteristics (52, 53). Generating a model using pressure data yields similar *in vivo* results compared to a steady-state velocity-based flow condition due to availability of time-resolved pressure data. Consequently, we used colonic pressure rather than fluid flow to develop the proposed model. However, it should be noted that colonic motility is also dependent on other factors, including faecal viscosity, which varies with diet, and understanding this relationship requires further study. However, it is important to note that there is limited research and specific data on the rheological properties of mucus and faeces. It is highly challenging to study these properties owing to various factors, including interpersonal differences, genetics, and rapid property changes when samples are removed from the body. The relaxation time (λ) and viscosity (µ) are some of key parameters for defining the viscoelastic behaviour of mucosal components in our model. These values are based on limited data available in the literature with minor adjustments (51, 54). To prevent mesh-related accuracy issues and possible convergence errors, we employed an unstructured mesh with characteristic size of 1.35 mm in the bulk and 0.1 mm near the crypt as shown in Fig. 2a. We studied several mesh resolutions, and we used the mesh values shown in Fig. 2a as the best compromise between accuracy and computational time.

**Figure 2.**
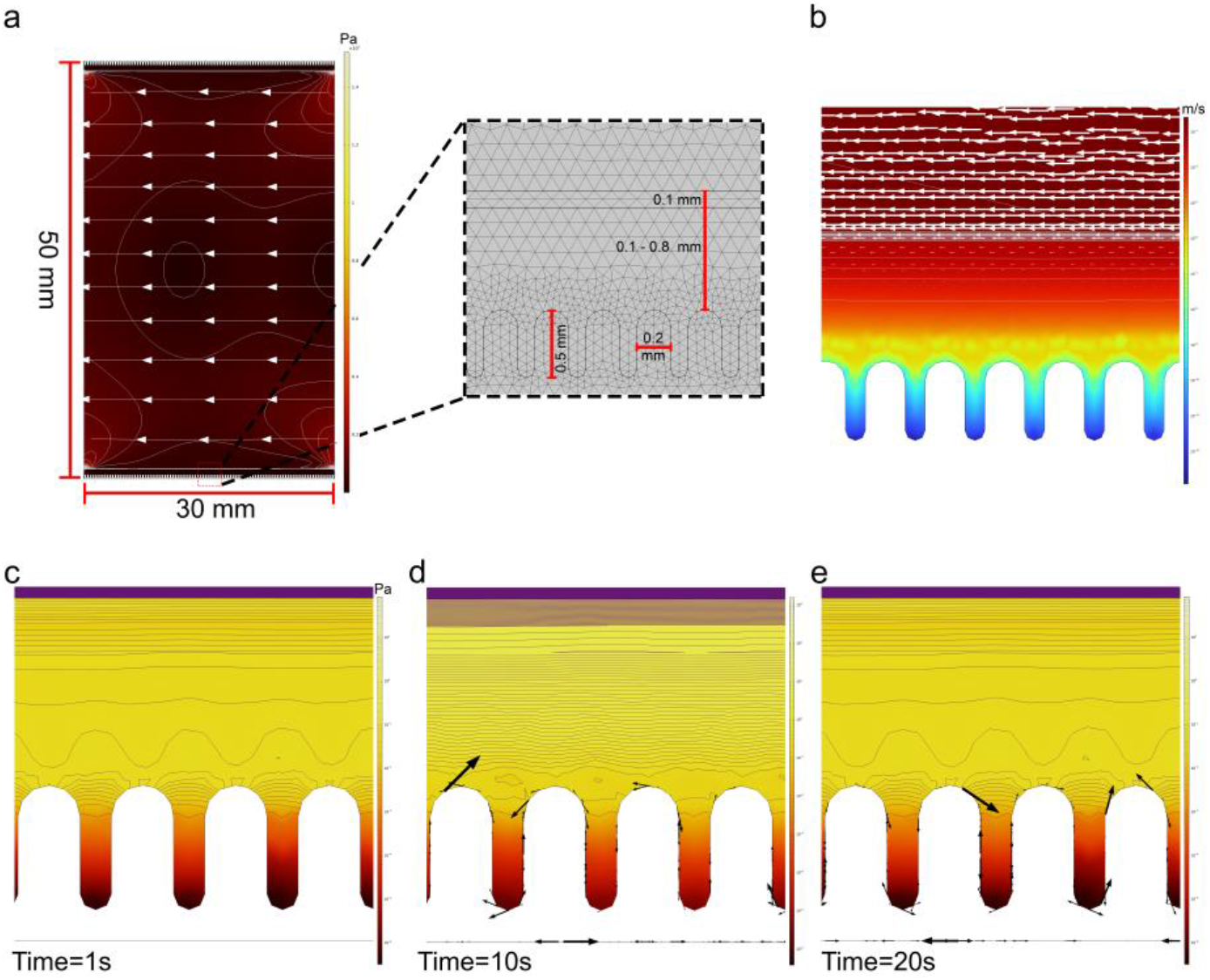
Homeostatic intestinal epithelium with 600 µm firmly attached mucus and demonstration of ternary phase field with fluid–structure interaction. (a) Flow-induced shear stress in the entire geometry is shown logarithmically with a colour bar (Pa), where the streamlines (white) indicate the velocity field, and the contour indicates the shear stress (Pa). The top and bottom parts of the system contained a mucus bilayer and embedded crypt structures. The thickness of the loosely attached mucus is 100 µm in all models, and the thickness of firmly attached mucus varies from 100 to 800 µm; (b) measured velocity magnitude at t=10s near crypt structures logarithmically with a colour bar (m/s), white arrows indicate the velocity field and grey lines indicate velocity magnitude (m/s). (c-d) Visualization of the shear stress distribution near crypt structures at t=1s, t10s, and t=20s. The contour in the Gaia colour range indicates the shear stress (Pa), and the black arrows indicate the shear component of traction on the crypt structures proportionally.

### 3.2 Mucus increases faecal matter velocity

Fluid flow due to pressure waves was simulated in the system, and the results were analyzed to observe the developing velocity field and total mass accumulated at the outlet. The velocity in the system shows a typical laminar flow with a low velocity near the walls and a peak velocity in the middle of the channel. The average Reynolds number in the mucosal models was 4.13^·^10^-6^ while in the UC model it was 5.23^·^10^-8^. Furthermore, in the UC model, luminal velocity in the channel centre point was 2.71^·^10^-5^ m/s at pressure wave peak compared to 1.4^·^10^-3^, 1.6^·^10^-3^, 2.3^·^10^-3^, 2.5^·^10^-3^, and 2.6^·^10^-3^ m/s for 100, 200, 400, 600, and 800 µm in the mucus models, respectively. On average, the mucus models had a velocity two orders of magnitude higher than that of the UC model (Fig. 4d). As the mucus layer acts as a lubricant to ease the movement of faeces in the *in vivo* colon, lack of mucus is likely to cause a reduction in this ability. In the mucosal models, velocity was significantly higher in the lumen above the mucus layers, indicating that mucus reduced velocity in the local area (Fig. 2b). On the other hand, in the UC model, the velocity continuously decreases towards crypt structures (Fig. 3b), supporting the hypothesis of mucus lubricating and therefore enhancing the velocity of faeces. The simulated net velocity value was within the range of the *in vivo* colonic content velocity value of 4.7^·^10^-3^ measured by MRI (48). Furthermore, in parallel with our results, the colonic content velocity in the human UC colon is often reduced (55, 56). Finally, we calculated the amount of accumulated feces at the outlet owing to the pressure-guided flow. The accumulated faecal matter volume are in agreement with the physiological levels; on average 18.3 g of matter was removed from the mucosal systems, whereas this value was 0.24 g in the UC model (Fig. 4e). Collectively, our numerical results suggest that colonic mucus may play a significant role in colonic content velocity and transport of faeces.

**Figure 3.**
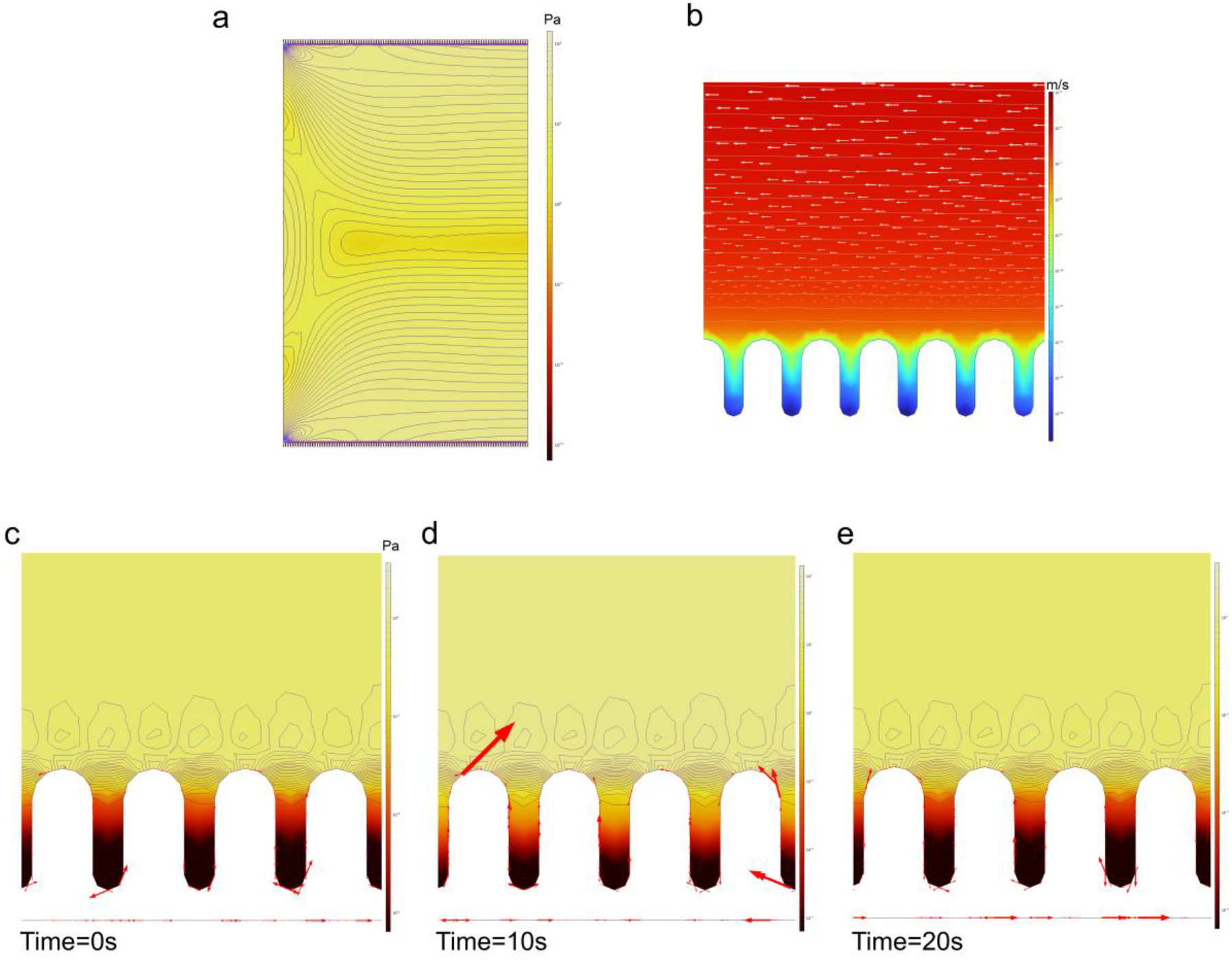
Velocity and shear stress distribution in UC epithelium without mucus. (a) Flow-induced shear stress in the entire geometry, contour shows the distribution in the model and the colour shows logarithmic shear values (Pa). (b) Measured velocity magnitude near crypt structures at t=10s, white arrows indicate the velocity field, colour shows logarithmic velocity values (m/s). (c-d) Visualization of shear stress distribution near crypt structures at t=1s, t10s, and t=20s respectively. The contour indicates the shear stress distribution, red arrows indicate the shear component of traction on crypt structures proportionally, colour indicates logarithmic values of shear (Pa).

**Figure 4.**
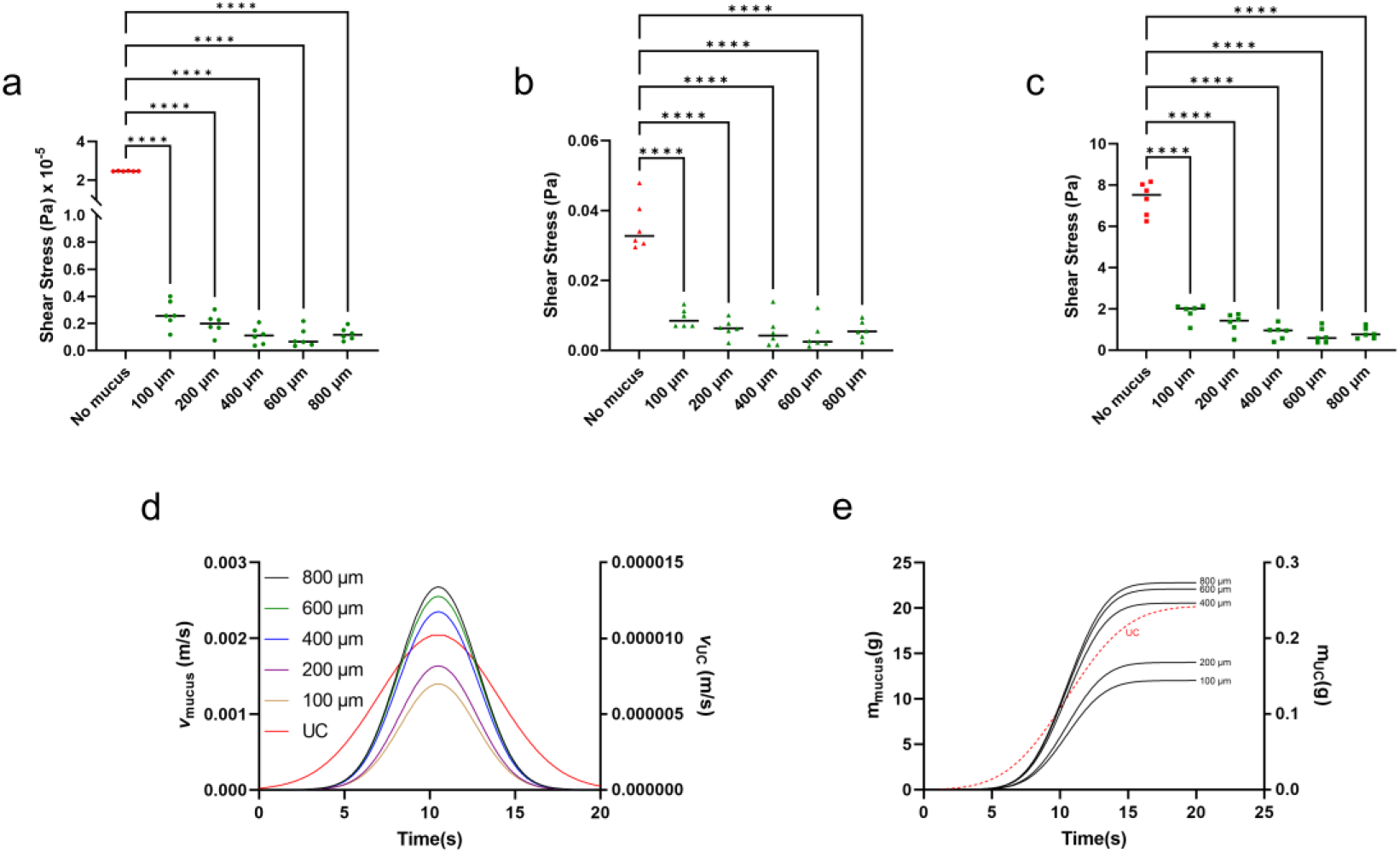
Statistical analysis of shear distribution on biologically important crypt sites: (a) crypt base, (b) crypt migration route, and (c) crypt top. Data are presented as the mean ±SD (n=6). ****P<0.0001. (d) average velocity magnitude (m/s) in the system, left y axis shows the values for mucus model while right y axis shows the values for UC model, (e) total faecal output (g) over time.

### 3.3. Mucus protects colonic crypts from motility induced shear

Current numerical models of the intestine focus on pharmacological aspects, such as transport (57), absorption, and drug release (58-60) while the effects of shear stress and mucus are often neglected. Although it is widely accepted that the mucus bilayer protects against mechanical forces, the effect of differences in the mucus bilayer, such as its thickness, or its complete absence in conditions such as IBD is not investigated. Overwhelming evidence suggests that shear stress has a significant impact on *in vitro* cellular behaviour, including structural organization (13), differentiation (61), and polarization (14) as well as microbiome behaviour through the regulation of mucus (62). Therefore, we theoretically investigated the spatial distribution of shear stress over time under physiological faecal flow conditions by performing a numerical analysis on a ternary-phase field model with an FSI composed of faeces, mucus bilayer, and colonic epithelium. To demonstrate the impact of shear stress, we mapped and compared the shear distribution of crypts in the UC and mucus models . Further, we varied the thickness of the SL mucus layer to mimic various pathological UC conditions where mucus thickness decreases depending on the severity of inflammation, and zones adjacent to the inflammatory site increase mucus production (32, 33). Our analysis indicated a crucial interplay between mucus thickness and shear stress delivered to crypt structures.

In the UC model, luminal shear in the channel centre point is 1.06 Pa after 10s, when the pressure wave peaks, while it is 6.35, 9.9, 13.9, 14.7, and 19.6 Pa in the mucus models with thickness 100, 200, 400, 600, and 800 µm, respectively. These results are consistent with the velocity results since higher velocities cause higher shear. Further statistical analysis of shear stress variance in the mucus (Fig.2c-e) and UC models (Fig.3c-e) near the crypt structures reveals significantly higher values in UC model without mucus at the crypt base (Fig. 4a), crypt migration route (Fig. 4b), and crypt top (Fig. 4c). These results show that our mucus models are consistent with observations indicating that the absence of mucus causes a significant increase in shear. Shear levels in the mucus models confirm that higher mucus thicknesses yield to relatively lower shear transfer to the crypt structures (Fig. 4a-c). Kotla et al. recently reported that hydrogel-based physical barrier placement significantly reduced inflammation in a mouse colitis model (63). Their results also suggest that a physical surface matrix coating to the inflamed colonic zones can be effective in reducing inflammation and restoration of the epithelial barrier function. Although the shear distribution in mice intestine is unclear, such a barrier with viscoelastic properties may play the role of mucus in shear dissipation, allowing mucosal healing.

In the crypt base, the UC model exhibits low variance in shear (SD =0.013) and is approximately 10 orders of magnitude higher than the mucus models. Among the mucus models, shear exhibits a negative trend, with 600 µm exhibiting the lowest values. Interestingly, 800 µm shows similar values to 400 µm at all three measurement sites. To better understand the effect of shear, we analyzed shear-induced tractions on crypts, which indicated that in both UC and mucus models, tractions over time exhibits behaviour that aligned with epithelial cell migration from the crypt base to the top.

Our model suggests that from the crypt base to the top, the average shear stress continuously increases from 0.094^·^10^-5^ to 2.47^·^10^-5^ Pa to 0.71 and 7.34 Pa in the homeostatic and UC models, respectively. These results, suggest that the mucus is a key component in the model to protect the underlying epithelium from the shear stress. Such differences in shear stress have been suggested to affect cellular behaviour (12). Our numerical results are consistent with the theory according to which shear drives intercellular forces resulting in cell migration dynamics from the crypt base to the top. Importantly, mechanical variables reportedly enable tissue folding, cell migration, and crypt morphogenesis in the intestinal organoids. Recently, Yang et al. reported that pressurization of the lumen drives crypt morphogenesis in intestinal organoids, while Pérez-González et al. reported that epithelial shape is driven by patterned forces that drive cell migration in intestinal organoids (21, 64). In agreement with our numerical analysis, they suggested an active cell crawling away from the crypt base, in which radial tractions point opposite to cell velocity. Finally, such an increase in shear from base to top may have a role in cellular processes such as differentiation and immune surveillance.

Across our mucus and UC models, the zone near crypt top has the highest shear stress values compared to other crypt sites. Further, the shear tractions were observed to start at the corners of the crypt top and merge at the middle position. Interestingly, this zone has been shown to be the site where cell-shedding occurs (20). Rapid renewal of the intestinal epithelium requires continuous cell shedding and replacement via cell migration. Dysregulation of cell shedding is linked to the development of IBD by creating gaps in the intestinal epithelium (65). Although the epithelial shedding process has been previously studied, little is known about the regulation of constitutive physiological shedding (20, 66). Recently, Marchiando et al. reported that tumour necrosis factor (TNF) increased epithelial shedding and apoptosis (67). Furthermore, shear stress is a key regulator of endothelial cell apoptosis (68). Several studies have also reported that high shear stress may promote the activation of immune cells, including T cells, neutrophils, and macrophages (15, 16, 69) and pro-inflammatory cytokine secretion(70). Therefore, this observation suggests that shear stress may have an important role in the cell-shedding process. Collectively, the high shear zone at the crypt top may cause pro-inflammatory activation, which can lead to epithelial apoptosis under homeostatic conditions as a contributor to cell-shedding processes. In UC, such high shear stress may play a role in UC pathogenesis through rapid differentiation of intestinal stem cells and/or altering cell migration dynamics.

Although our study is limited to 2D simulations, the result show for the first time the time-space dynamics of shear stress on colonic epithelium and how mucus affects this process. The model predictions are consistent with *in vitro* observations: UC manifestation significantly affects colonic mucus thickness, and a physical barrier on inflamed zones can significantly lower epithelial damage. Our findings indicate that there is a significant interplay between velocity, shear stress, and mucus. All of which may play an essential role in both intestinal inflammation and homeostasis. Shifting the focus to mucus and shear stress might be the key to achieving better microphysiological models that could provide unique mechanistic insights in homeostasis and inflammation. Combining our model with other computational models may have implications for the development of novel therapeutics in terms of physical interactions and may have wide use in understanding absorption, drug retention time, particle size optimization, and compound/surface interactions.

## Acknowledgments

This work was supported by Science Foundation Ireland 18/EPSRC-CDT/3583 and the Engineering and Physical Sciences Research Council EP/S02347X/1. This work has emanated from research supported in part by a grant from Science Foundation Ireland (SFI) and the European Regional Development Fund (ERDF) under grant number 13/RC/2073_P2

